# Forearm and hand muscles exhibit high coactivation and overlapping of cortical motor representations

**DOI:** 10.1101/2021.07.20.453121

**Authors:** Gabriela P. Tardelli, Victor Hugo Souza, Renan H. Matsuda, Marco A. C. Garcia, Pavel A. Novikov, Maria A. Nazarova, Oswaldo Baffa

## Abstract

Most of the motor mapping procedures using navigated transcranial magnetic stimulation (nTMS) follow the conventional somatotopic organization of the primary motor cortex (M1) by assessing the representation of a particular target muscle, disregarding the possible coactivation of synergistic muscles. In turn, multiple reports describe a functional organization of the M1 with an overlapping among motor representations acting together to execute movements. In this context, the overlap degree among cortical representations of synergistic hand and forearm muscles remains an open question. This study aimed to evaluate the muscle coactivation and representation overlapping common to the grasping movement and its dependence on the stimulation parameters. The nTMS motor maps were obtained from one carpal muscle and two intrinsic hand muscles during rest. We quantified the overlapping motor maps in size (area and volume overlap degree) and topography (similarity and centroid’s Euclidean distance) parameters. We demonstrated that these muscle representations are highly overlapped and similar in shape. The overlap degrees involving the forearm muscle were significantly higher than only among the intrinsic hand muscles. Moreover, the stimulation intensity had a stronger effect on the size compared to the topography parameters. Our study contributes to a more detailed cortical motor representation towards a synergistic, functional arrangement of M1. Understanding the muscle group coactivation may provide more accurate motor maps when delineating the eloquent brain tissue during pre-surgical planning.

## Introduction

A fundamental debate on primary motor cortex (M1) organization is whether different body parts rely on a discrete somatotopic or functionally-specific representation on the cortical surface (Schieber 2001). In the M1, the somatotopic organization associates a cortical site to the control of a specific muscle (Penfield and Rasmussen 1950), whereas the functional organization suggests the cortical representation of limb movements (Gentner et al. 2010; Strother et al. 2012). Several studies have demonstrated that the high complexity of central movement generation can be derived from an extensive overlap and redundancy between adjacent cortical area representations (Schieber 2001; Devanne et al. 2006; Gentner and Classen 2006; Melgari et al. 2008). The overlapping areas can be related to specific movements involving more than one adjacent single joint and, therefore, a complex synergy among different muscles (Strother et al. 2012; Leo et al. 2016), which corroborates a hypothesis of the functional organization of M1. In this context, the overlap degree (OD) in the cortical representation of synergistic hand and forearm muscles remains an open question.

In navigated transcranial magnetic stimulation (nTMS), a coil placed on the scalp over M1 produces magnetic pulses that induce electric fields in the cortical tissue. The neuronal excitation results in action potentials that propagate through the corticospinal tract generating motor evoked potentials (MEP). The MEP amplitude combined with the TMS coil coordinates of the individual’s brain enables one to delimit the extension and location of the motor cortical representations of the body parts (Romero et al. 2011). The nTMS mapping is widely used for delineating eloquent motor function in a preoperative setting (Lefaucheur and Picht 2016; Krieg et al. 2017). An approach that accounts for the functional overlap in cortical motor representations can lead to more selective cortical maps with the potential to improve patient prognostics (Frey et al. 2014; Picht et al. 2016).

A few studies claim that the overlap in cortical representation may partially represent a cortical manifestation for synergies (Pearce et al. 2000; Tyc and Boyadjian 2005; Latash et al. 2007; Cheung et al. 2012; Overduin et al. 2012; Leo et al. 2016; Huffmaster et al. 2018; Raffin and Siebner 2019). In this model, multiple muscle groups form different synergy patterns producing complex movements (Fricke et al. 2020). To the best of our knowledge, most conventional motor mapping procedures assess the cortical representation of a particular target muscle (Krieg et al. 2017), disregarding the possible synergistic activation of the adjacent muscles. This coactivation of muscle groups may provide further information about how and to what degree synergistic muscles are represented at the cortical level (Leo et al. 2016).

The aim of our study was to quantify the OD between the cortical motor representation of two intrinsic hand muscles and one carpal forearm muscle in rest conditions. Using nTMS mapping, we delineated the motor representation of the selected muscles considering their synergistic activation. We hypothesized that these representations would be highly overlapped due to the muscles’ extensive coactivation in several hand movements, e.g., grasping. Also, the OD would differ between adjacent and target muscles from different body parts and would increase with the TMS intensity. Our results provide novel evidence on the functional cortical motor organization of the human brain.

## Material and Methods

### Participants

The experiment was performed with 12 right-handed (Edinburgh handedness inventory (Oldfield 1971), mean score: +75; range: +55 to +95) young male volunteers (mean age: 31.3 ± 2.5 years; range: 27–35 years). Participants were asymptomatic to neurological and psychiatric disorders, without recurrent headaches, and free of medication during the data collection phase. The experimental procedure followed the Declaration of Helsinki and was approved by the local ethical committee (CAAE: 54674416.9.0000.5407). Before the testing procedures, all participants signed a consent form.

### Experimental procedure

Subjects underwent a magnetic resonance imaging (MRI) scan (Achieva 3T; Philips Healthcare, Netherlands) with a T1-weighted gradient-echo sequence (acquisition matrix 240×240×240, voxel size 1×1×1 mm^3^, 6.7 ms repetition time, and 3.1 ms echo time). The gray matter surface of the brain was segmented using SPM 12 software (Friston et al. 2006) for guiding the nTMS coil placement. Surface EMG electrodes (circular 10-mm diameter; model 2223 BRQ, 3M Brazil Ltd., Sumaré, Brazil) were placed in a pseudo-monopolar montage, with one electrode over the innervation zone and the other over the closest bone eminence (Garcia et al. 2017, 2020). The selected muscles were one right carpal forearm muscle (*flexor carpi radialis;* FCR), and two right intrinsic hand muscles, a thenar (*flexor pollicis brevis;* FPB) and a hypothenar muscle (*abductor digiti minimi;* ADM). EMG data were continuously recorded from the three muscles and digitized with the EMG 410C amplifier (gain: 2000 x, sampling frequency: 3.5 kHz per channel, band-pass 4^th^-order Butterworth filter: 20-500 Hz, A/D converter: 12 bits; EMG System do Brasil, São José dos Campos, Brazil).

The participants sat in a reclining chair and were instructed to stay fully relaxed with their right hand in a neutral posture during the nTMS session. TMS biphasic pulses were delivered with a figure-of-eight coil (10 cm diameter windings) connected to a Neuro-MS stimulator (Neurosoft, Ivanovo, Russia). The coil placement was guided by the neuronavigation software InVesalius Navigator (Souza et al. 2018) connected to the MicronTracker Sx60 (ClaroNav, Toronto, Canada) spatial tracker. Figure 1A depicts the experimental setup. The following procedure was applied separately for each muscle (FCR, FPB, and ADM). First, the hotspot was defined as the coil location showing the highest MEP amplitudes with the coil tangential to the scalp and approximately perpendicular to the central sulcus (Bashir et al. 2013; Souza et al. 2017). Second, the resting motor threshold (rMT) was defined as the minimum stimulator intensity at which the MEP amplitudes were greater than 100 μV in 5 out of 10 pulses (Nielsen 1996; Nogueira-Campos et al. 2014). A higher threshold amplitude than the usual 50 μV (Conforto et al. 2004) was selected to provide stable MEP measurements desired when using a relatively small number of trials per stimulation site during motor mapping (Pellegrini et al. 2018).

**Fig. 1.**
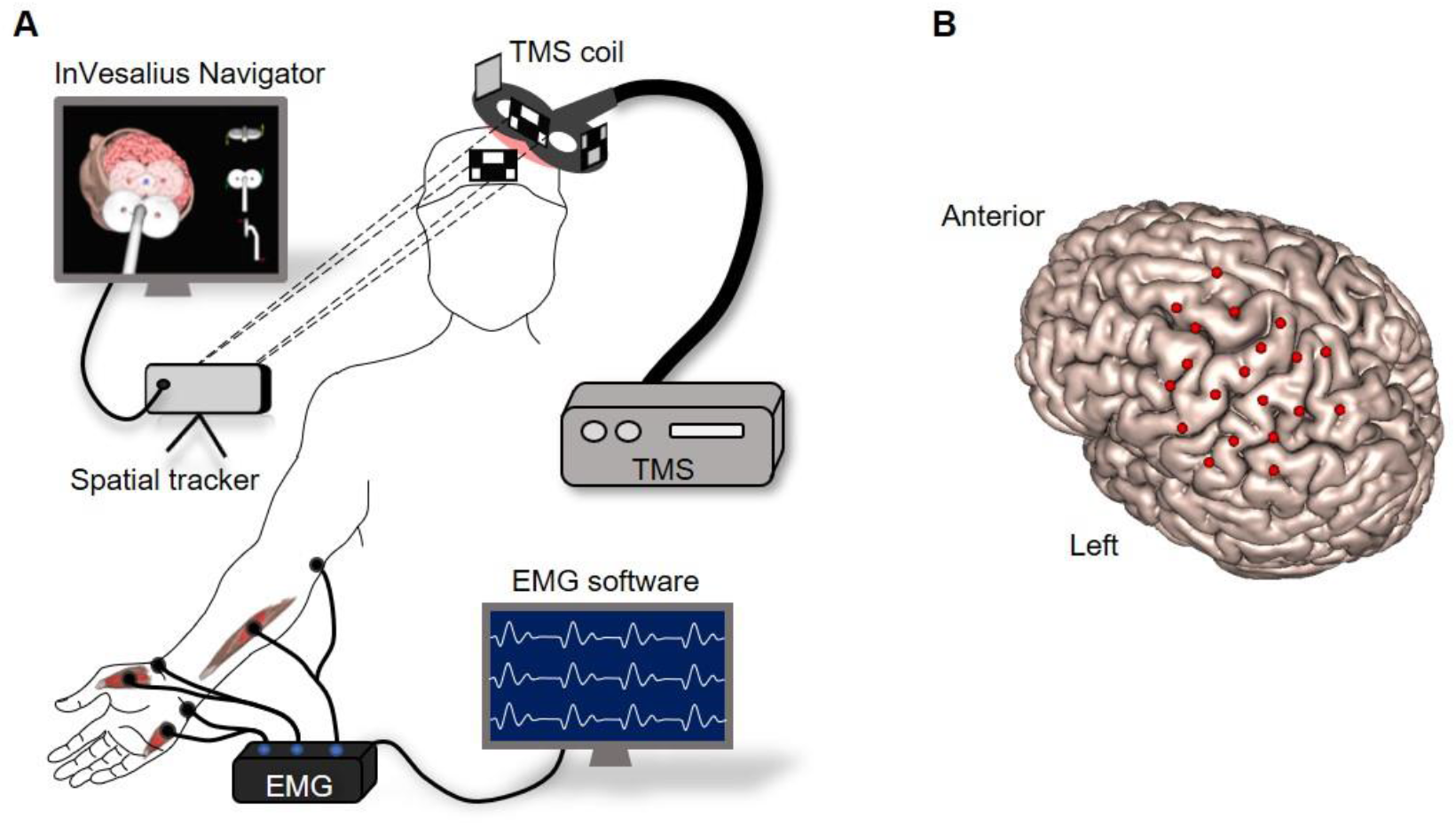
A) Schematic representation of the experimental setup. The MEPs were recorded simultaneously from the hand and forearm muscles while TMS was applied over the left hemisphere guided by the InVesalius Navigator software. B) Grid of coil center locations relative to the cortical surface. This motor map was recorded at a stimulation intensity of 120% of rMT of the FCR muscle on a representative subject. The grid was centered on the muscle’s hotspot, and each coil location is the average across three trials.

Motor mapping was performed following the pseudo-random walk method with three consecutive TMS pulses in each of the 20 sites around the target muscle’s hotspot (Van De Ruit et al. 2015; Cavaleri et al. 2018; Jonker et al. 2018). Each motor map composed an unevenly spaced grid centered on the hotspot for each individual, as illustrated for a representative subject in Fig 1B. The distance between adjacent stimulation sites was approximately 14.5 ± 3.6 mm, and interpulse intervals were pseudo-randomized between 5 and 10 s. EMG signal was recorded from the three muscles simultaneously, and the coordinates of the coil center were recorded with the neuronavigation software. A trigger signal synchronized the EMG and the neuronavigation software. The target muscle was defined as the one whose stimulation intensity was set relative to its rMT, and the stimuli were delivered over a region centered at the hotspot. The remaining muscles were defined as adjacent. The experiment was repeated for each target muscle (ADM, FCR, and FPB) and with stimulation intensities of 110% and 120% of the rMT, scaled according to the maximum stimulator output (MSO).

### Motor map processing

The EMG signals were processed using the Signal Hunter software (https://doi.org/10.5281/zenodo.1326308) written in Matlab R2017a (MathWorks Inc., Natick, USA). The peak-to-peak amplitude was computed for MEPs extracted from the EMG signal in a time window 10–60 ms after the TMS pulse. The EMG signal was visually inspected, and trials with muscle pre-activation, artifacts, or noise over ± 20 μV up to 300 ms before the TMS pulse in amplitude were rejected. After the preprocessing, all subjects had three trials in each of the 20 cortical targets per motor map, except for one subject in which three out of the 60 stimulations were rejected due to muscle pre-activation and noisy EMG signals. The coil center coordinates obtained from InVesalius Navigator were imported into Signal Hunter and aligned to the corresponding MEP amplitude and the latency of the three muscles. One participant was removed from the data analysis due to technical problems in the TMS–EMG synchronization.

The cortical motor maps were created in the TMSmap software (Novikov et al. 2018) with the individuals’ MRI, stimulation coordinates, and the correspondent MEP amplitudes. The software creates the maps with the mean coordinates and the median peak-to-peak MEP amplitudes of the merged closely spaced coordinates, resulting in a cortical motor map with 20 MEP amplitudes and coil locations per tested condition for each subject. The technical details of the map processing steps performed in the TMSmap software are described in Appendix 1. For each target muscle, we constructed two overlaps: the target with each of the adjacent muscles (two maps) and the target with both adjacent muscles together (three maps). The area, volume, and centroid were computed for all maps from each subject and stimulation intensity. The area represents the extent of the cortical motor representation, and the volume represents the area weighted by the motor response amplitude. To quantify the topographic similarity between two (or three) maps, we computed the Earth’s movers distance (EMD), which estimates the work required to move one spatial distribution to another (Rubner et al. 1998). The Euclidean distance between the target muscle and the overlap map centroids of the corresponding target and its adjacent muscles was computed to evaluate differences in the spatial distribution of the muscle of interest when overlapped with another map. We defined the EMD and the Euclidean distance as topography parameters.

To evaluate the coactivation between the muscle representations, we defined the size parameters as area and volume OD. The OD was computed as the percentage of the area (or volume) that evoked two or three muscles relative to the total area (or volume) that evoked at least one of the assessed muscles (Melgari et al. 2008; Nazarova et al. 2021):

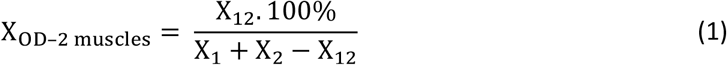

Where X can be area or volume, and the indices 1, 2, and 3 (equation below) refer to each of the muscle maps (target and adjacent) and their corresponding overlaps (pairs of indices). The OD was categorized as: 0-20% (negligible); 21-40% (low); 41-60% (medium); 61-80% (high); 81-100% (very high). The relative number of subjects with OD in each of these categories was calculated for all overlap maps. Similarly, the OD of three muscle motor maps were computed as:

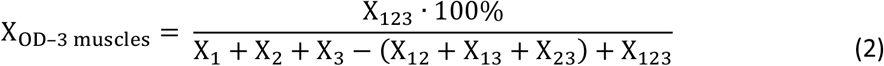

### Statistical analysis

All parameters were normalized relative to their maximum values within each individual to enable a direct comparison between conditions and subjects. The stimulation intensity, target, and adjacent muscle and each map parameter (area and volume OD, EMD, and the centroid Euclidean distance) were modeled as fixed effects. In turn, the subjects were modeled as a random effect in a linear mixed-effects model. A random structure of the model was selected based on hierarchical sequential testing with each model fit using likelihood-ratio tests. The chosen model was recomputed using restricted maximum likelihood estimation and the p-values for fixed effects derived with Satterthwaite approximations in a Type III Analysis of Variance table. When appropriate, posthoc multiple comparisons were performed with estimated marginal means with p-value correction for the false discovery rate. The rMT across subjects (random) and muscles (fixed) were also analyzed using a linear mixed-effects model, and multiple comparisons were performed using the Tukey simultaneous tests for the difference of means. Critical deviations from normality were assessed with the residuals’ Q–Q plots, and homoscedasticity was inspected with a standard versus fitted values plot. Statistical analysis was performed in R 3.6 (R Core Team, Vienna, Austria) using the *lme4* 1.1, *afex* 0.25 packages, and *emmeans* 1.4 packages. The level of statistical significance was set at 0.05.

## Results

### Overlap degree and rMT

The rMT varied across muscles (*p* = 0.039) and within subjects (*p* = 0.015), being higher for the FCR compared to the FPB muscle (*p* = 0.042) and similar when comparing ADM with FCR (*p* = 0.120) or FPB (*p* = 0.852) muscles. We computed nine overlap maps for each subject: six overlap maps of two muscles and three overlaps of three muscles. The subscript [*tg*] refers to the target and [*adj*] to the adjacent muscles. The relative number of subjects for each OD category is illustrated in Figure 2. More than 60% of the subjects had medium to very high area OD but negligible to low volume OD. Maps of two muscles had more subjects with higher ODs than those of three muscles at both stimulation intensities. The number of subjects with high area OD of all maps was higher at 120% than at 110% rMT stimulation intensity.

**Fig. 2.**
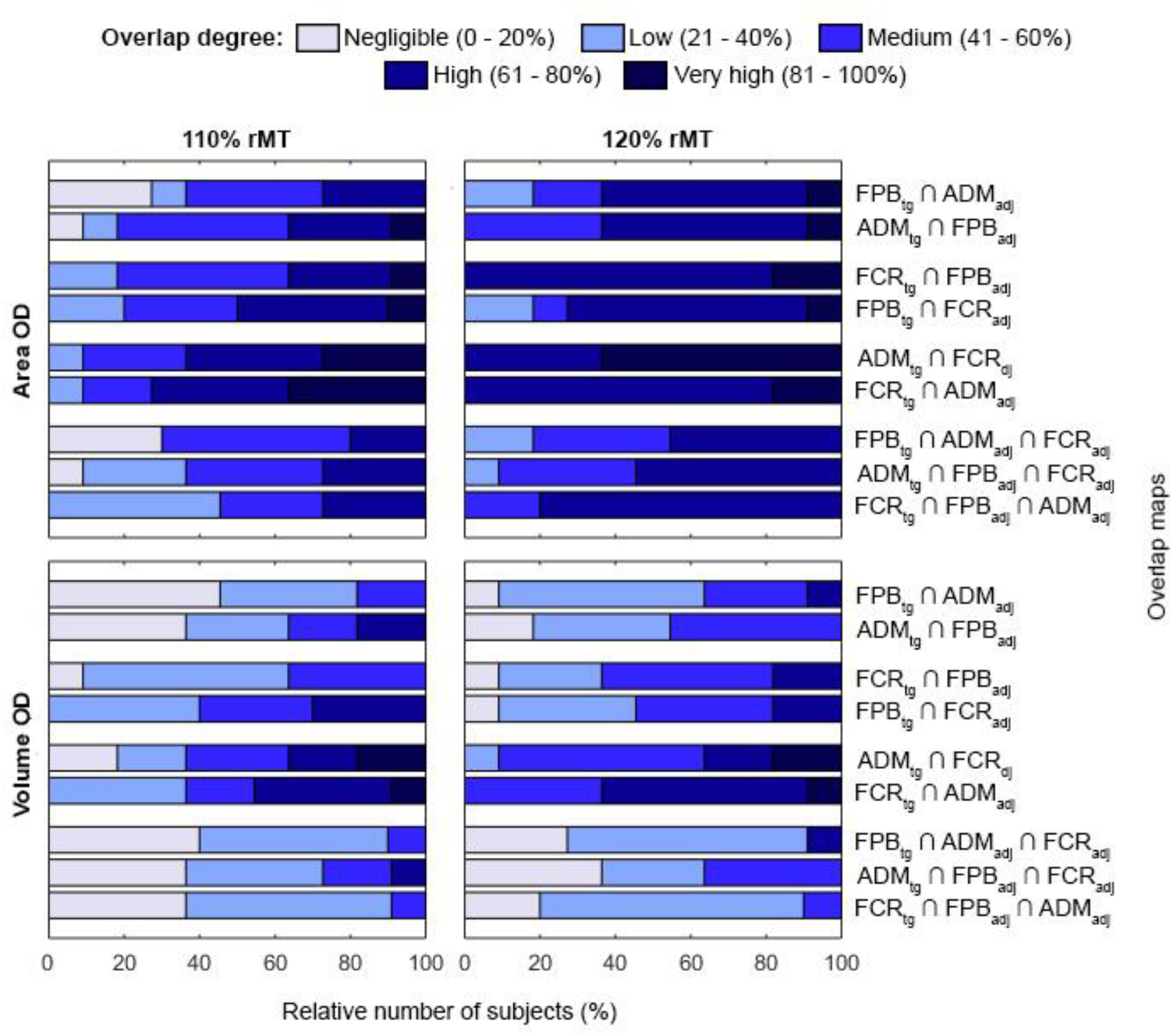
Relative number of subjects with OD distributed in 5 categories (negligible, low, medium, high and very high) for all overlap maps and at stimulation intensities of 110% (left) and 120% (right) of rMT. The length of the horizontal bars indicates the relative number of subjects with each OD, and the ⋂ (intersection) symbol represents the overlap between muscles.

### The effect of stimulation intensity, target and adjacent muscle on map size and topography

Table 1 presents the results of the linear mixed-effects model. The size parameters (area and volume OD) were significantly affected by the stimulation intensity, the target and adjacent muscle individually, and their interaction. In turn, the topography parameters (EMD and the centroid Euclidean distance) were significantly affected only by the adjacent muscle but not by the stimulation intensity nor the target muscle. The interaction between stimulation intensity and target muscle affected both topography parameters, while the interaction between target and adjacent muscles affected only the EMD.

**Table 1.** The linear mixed-effects model results for size (area and volume OD) and topography (EMD and centroid Euclidean distance) parameters. Abbreviations: degrees of freedom (DoF), stimulation intensity (SI), target (M_tg_), and adjacent muscle (M_adj_). The * indicates *p*-value < 0.05

| Effect | DoF<br>(numerator,<br>denominator) | F-value | $p$-value | DoF<br>(numerator,<br>denominator) | F-value | $p$-value |
| --- | --- | --- | --- | --- | --- | --- |
| Area OD |  |  |  | Volume OD |  |  |
| SI | (1, 10.0) | 19.99 | 0.001* | (1, 9.96) | 5.85 | 0.036* |
| $M_{tg}$ | (2, 10.7) | 8.61 | 0.006* | (2, 10.9) | 11.82 | 0.002* |
| $M_{adj}$ | (3, 137.4) | 45.77 | <0.001* | (3, 136.0) | 70.35 | <0.001* |
| SI x $M_{tg}$ | (2, 137.9) | 0.82 | 0.444 | (2, 136.6) | 0.06 | 0.945 |
| SI x $M_{adj}$ | (3, 137.4) | 0.09 | 0.964 | (3, 135.9) | 1.11 | 0.348 |
| $M_{tg}$ x $M_{adj}$ | (3, 137.4) | 7.02 | <0.001* | (3, 136.0) | 13.01 | 0.000* |
| SI x $M_{tg}$ x $M_{adj}$ | (3, 137.5) | 1.30 | 0.276 | (3, 136.0) | 0.70 | 0.553 |
| Centroid Euclidean distance |  |  |  | EMD |  |  |
| SI | (1, 11.0) | 0.02 | 0.890 | (1, 10.1) | 1.65 | 0.228 |
| $M_{tg}$ | (2, 11.1) | 0.39 | 0.688 | (2, 14.5) | 2.76 | 0.096 |
| $M_{adj}$ | (3, 146.9) | 4.49 | 0.005* | (3, 147.5) | 5.57 | 0.001* |
| SI x $M_{tg}$ | (2, 147.1) | 4.06 | 0.019* | (2, 147.9) | 3.16 | 0.045* |
| SI x $M_{adj}$ | (3, 146.9) | 0.31 | 0.819 | (3, 147.4) | 0.28 | 0.836 |
| $M_{tg}$ x $M_{adj}$ | (3, 146.9) | 1.32 | 0.270 | (3, 147.5) | 4.82 | 0.003* |
| SI x $M_{tg}$ x $M_{adj}$ | (3, 146.9) | 1.41 | 0.241 | (3, 147.6) | 0.55 | 0.646 |

The multiple comparisons are presented in Tables 2–4, and the means and standard deviations across subjects for each parameter of all overlap maps area are illustrated in Figure 3. For most overlaps, the area OD was significantly higher at 120% than at 110% rMT of stimulation intensity. The intensity effect was only significant at the volume OD factor level, whereas none of the multiple comparisons displayed any significant differences. When comparing different target or adjacent muscles, the highest area and volume ODs were between ADM and FCR. In turn, overlaps involving FPB did not show any significant differences when compared to the overlap between the three muscles together. This result applies to both stimulation intensities in most cases. Lastly, the EMD and centroid Euclidean distance were similar for both tested stimulation intensities and all comparisons across adjacent and target muscles.

**Fig. 3.**
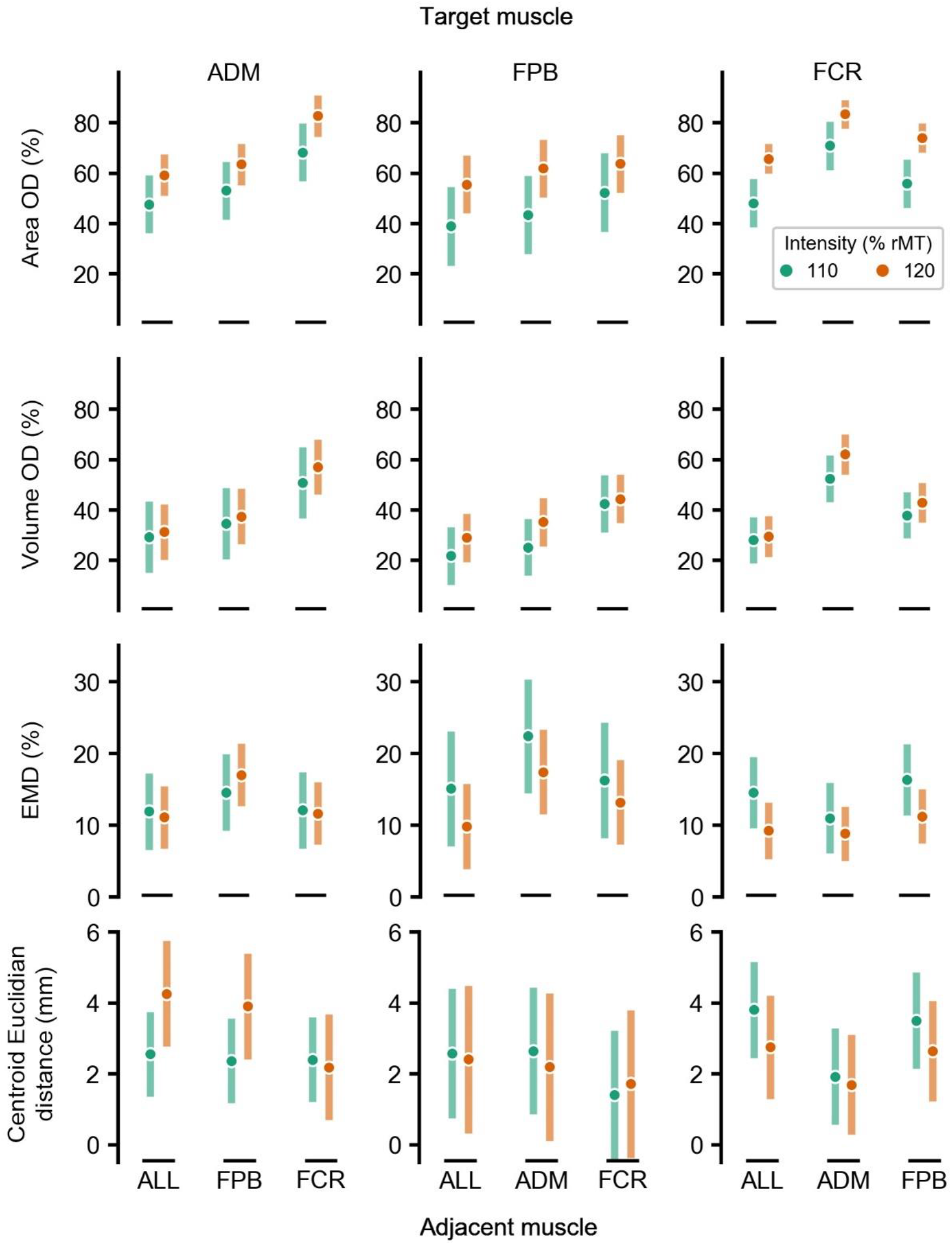
Mean and standard deviation of the motor maps’ size and topography parameters across the 11 subjects. The rows and columns of the chart grid contain the map parameters and the target muscle, respectively. The green and orange colors represent 110% and 120% of the rMT stimulation intensity. The term ALL refers to the overlap of the target with both adjacent muscles simultaneously

**Table 2.**
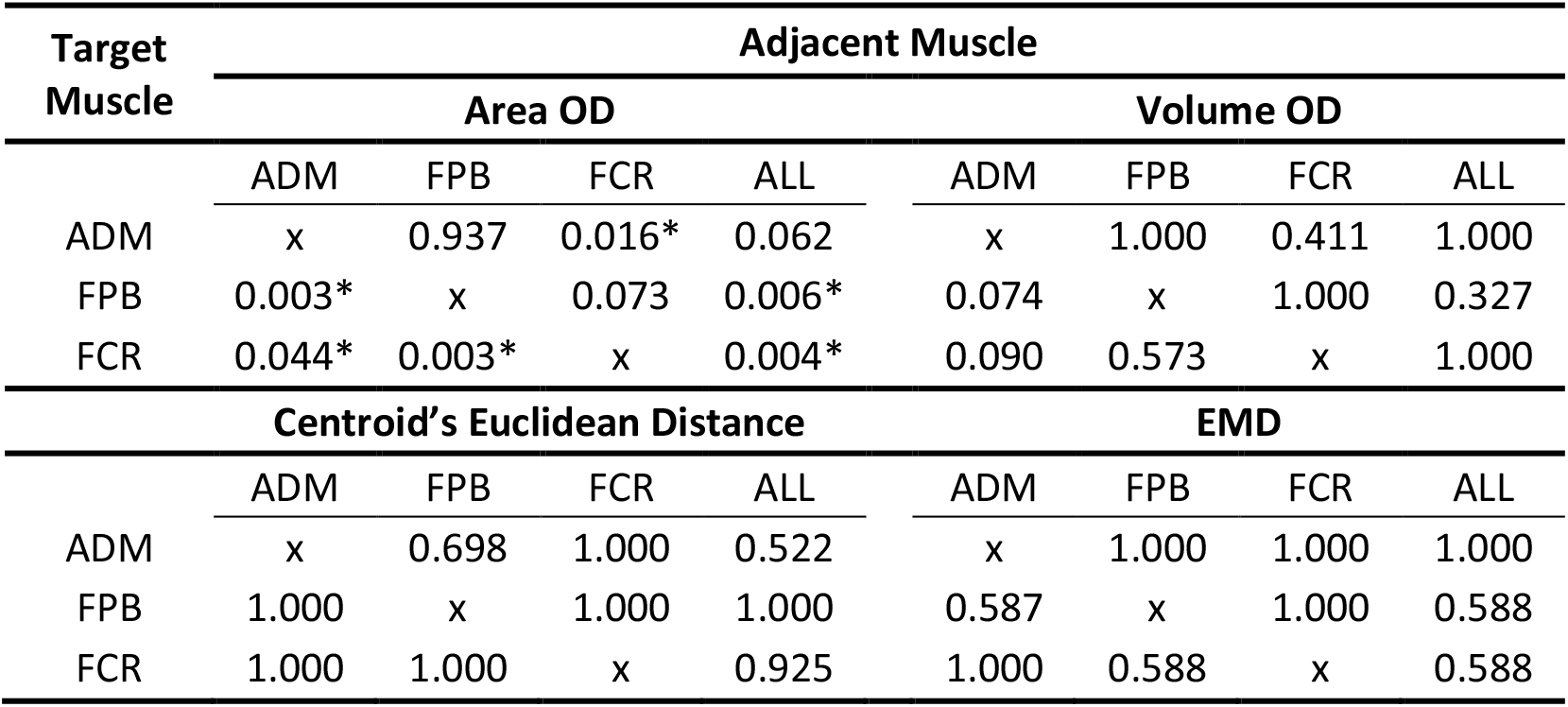
Multiple comparisons of area OD, volume OD, centroid’s Euclidean distance, and EMD between the stimulation intensities (110% compared with 120% of rMT) for each combination of target and adjacent muscles.

**Table 3.** Multiple comparisons of area OD, volume OD, centroid’s Euclidean distance, and EMD between different target muscles for each adjacent muscle and stimulation intensity (% of rMT).

| Muscles |  |  | Area OD |  | Volume OD |  | Centroid's Euclidean distance |  | EMD |  |
| --- | --- | --- | --- | --- | --- | --- | --- | --- | --- | --- |
| Intensity (% rMT) |  |  |  |  |  |  |  |  |  |  |
| Target 1 | Target 2 | Adjacent | 110 | 120 | 110 | 120 | 110 | 120 | 110 | 120 |
| FPB | FCR | ADM | 0.001* | 0.006* | <0.001* | <0.001* | 1.000 | 1.000 | 0.053 | 0.157 |
| ADM | FCR | FPB | 1.000 | 0.102 | 1.000 | 0.662 | 0.878 | 0.826 | 1.000 | 0.253 |
| ADM | FPB | FCR | 0.011* | 0.003* | 0.444 | 0.115 | 1.000 | 1.000 | 0.697 | 1.000 |
| ADM | FPB | ALL | 0.212 | 0.890 | 0.513 | 1.000 | 1.000 | 0.826 | 1.000 | 1.000 |
| ADM | FCR | ALL | 1.000 | 0.434 | 1.000 | 1.000 | 0.826 | 0.801 | 1.000 | 1.000 |
| FPB | FCR | ALL | 0.307 | 0.230 | 0.513 | 1.000 | 1.000 | 1.000 | 1.000 | 1.000 |

**Table 4.**
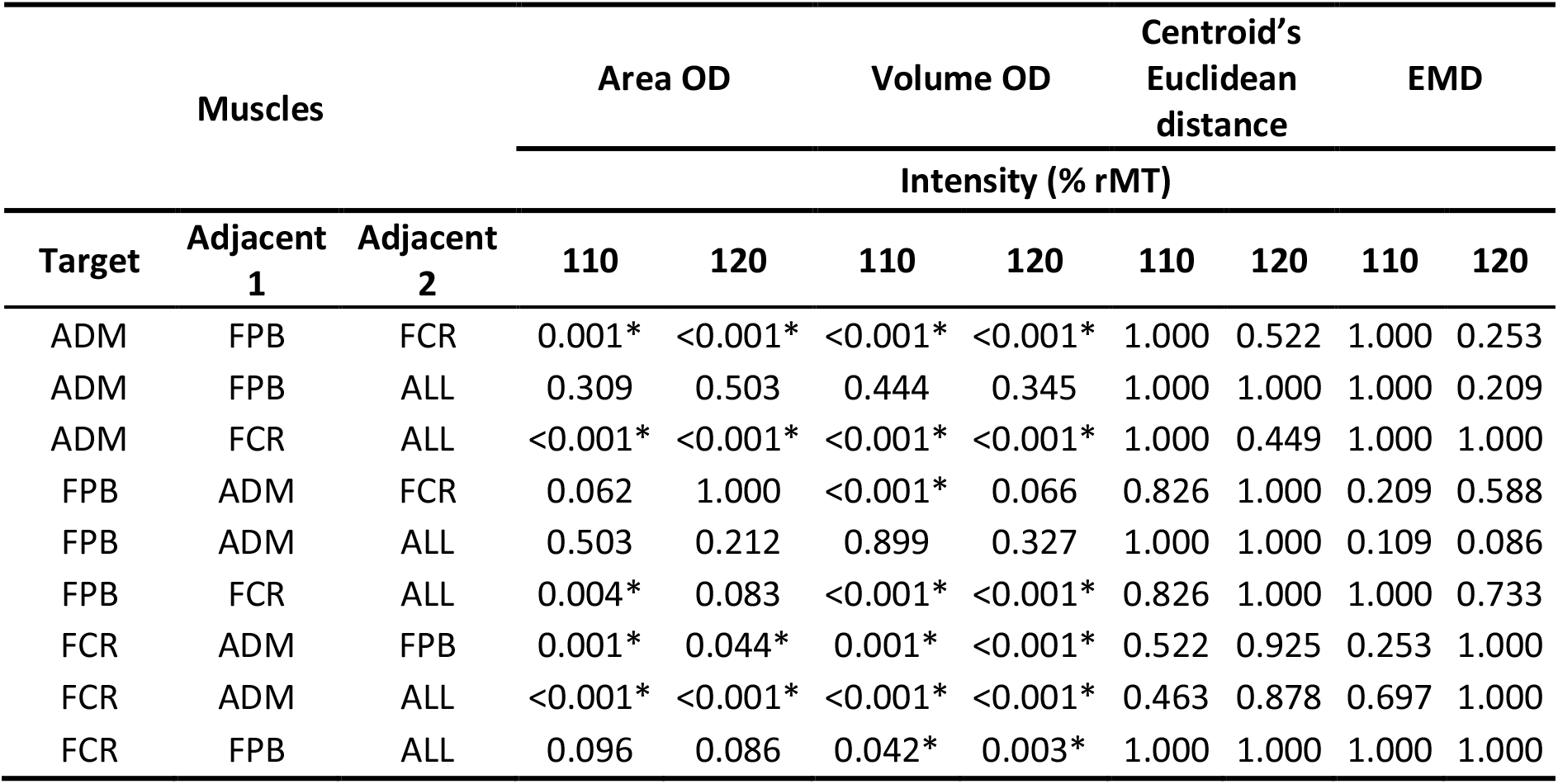
Multiple comparisons of area OD, volume OD, centroid’s Euclidean distance, and EMD between different adjacent muscles for each target muscle and stimulation intensity (% of rMT).

| Muscles |  |  | Area OD |  | Volume OD |  | Centroid's Euclidean distance |  | EMD |  |
| --- | --- | --- | --- | --- | --- | --- | --- | --- | --- | --- |
|  |  |  |  |  |  |  | Intensity (% rMT) |  |  |  |
| Target | Adjacent 1 | Adjacent 2 | 110 | 120 | 110 | 120 | 110 | 120 | 110 | 120 |
| ADM | FPB | FCR | 0.001* | <0.001* | <0.001* | <0.001* | 1.000 | 0.522 | 1.000 | 0.253 |
| ADM | FPB | ALL | 0.309 | 0.503 | 0.444 | 0.345 | 1.000 | 1.000 | 1.000 | 0.209 |
| ADM | FCR | ALL | <0.001* | <0.001* | <0.001* | <0.001* | 1.000 | 0.449 | 1.000 | 1.000 |
| FPB | ADM | FCR | 0.062 | 1.000 | <0.001* | 0.066 | 0.826 | 1.000 | 0.209 | 0.588 |
| FPB | ADM | ALL | 0.503 | 0.212 | 0.899 | 0.327 | 1.000 | 1.000 | 0.109 | 0.086 |
| FPB | FCR | ALL | 0.004* | 0.083 | <0.001* | <0.001* | 0.826 | 1.000 | 1.000 | 0.733 |
| FCR | ADM | FPB | 0.001* | 0.044* | 0.001* | <0.001* | 0.522 | 0.925 | 0.253 | 1.000 |
| FCR | ADM | ALL | <0.001* | <0.001* | <0.001* | <0.001* | 0.463 | 0.878 | 0.697 | 1.000 |
| FCR | FPB | ALL | 0.096 | 0.086 | 0.042* | 0.003* | 1.000 | 1.000 | 1.000 | 1.000 |

## Discussion

In this study, we demonstrate that the motor map parameters vary significantly between the pairs of muscles in size (area and volume) but not in topography (EMD and centroid Euclidean distance). While area OD of the muscle’s pairs was mainly medium to very high, volume OD was negligible to low. Our results also show that increasing the stimulation intensity from 110% to 120% of the rMT causes a significant increase in the area OD among different muscles. In turn, changing the stimulation intensity does not affect the map topographies. The volume OD does not seem to increase with the stimulus intensity, possibly due to OD variations between subjects and muscles.

### The effect of TMS intensity on the map parameters

Lower stimulation intensities resulted in a motor representation with more restricted sites of muscle activation than higher intensities, as depicted in Figure 4. One likely interpretation is that the weaker intensity might not recruit the less excitable neuronal populations (Kallioniemi and Julkunen 2016). When studying the functional overlapping of muscles with different motor thresholds, one might observe that M1 is organized by discrete or slightly overlapping representations. In each M1 representation site, the target muscle can be overlapped with different adjacent muscles that may be related to distinct synergies and, therefore, contribute to various movements. This association might point towards the functional organization of M1 (Massé-Alarie et al. 2017).

**Fig. 4.**
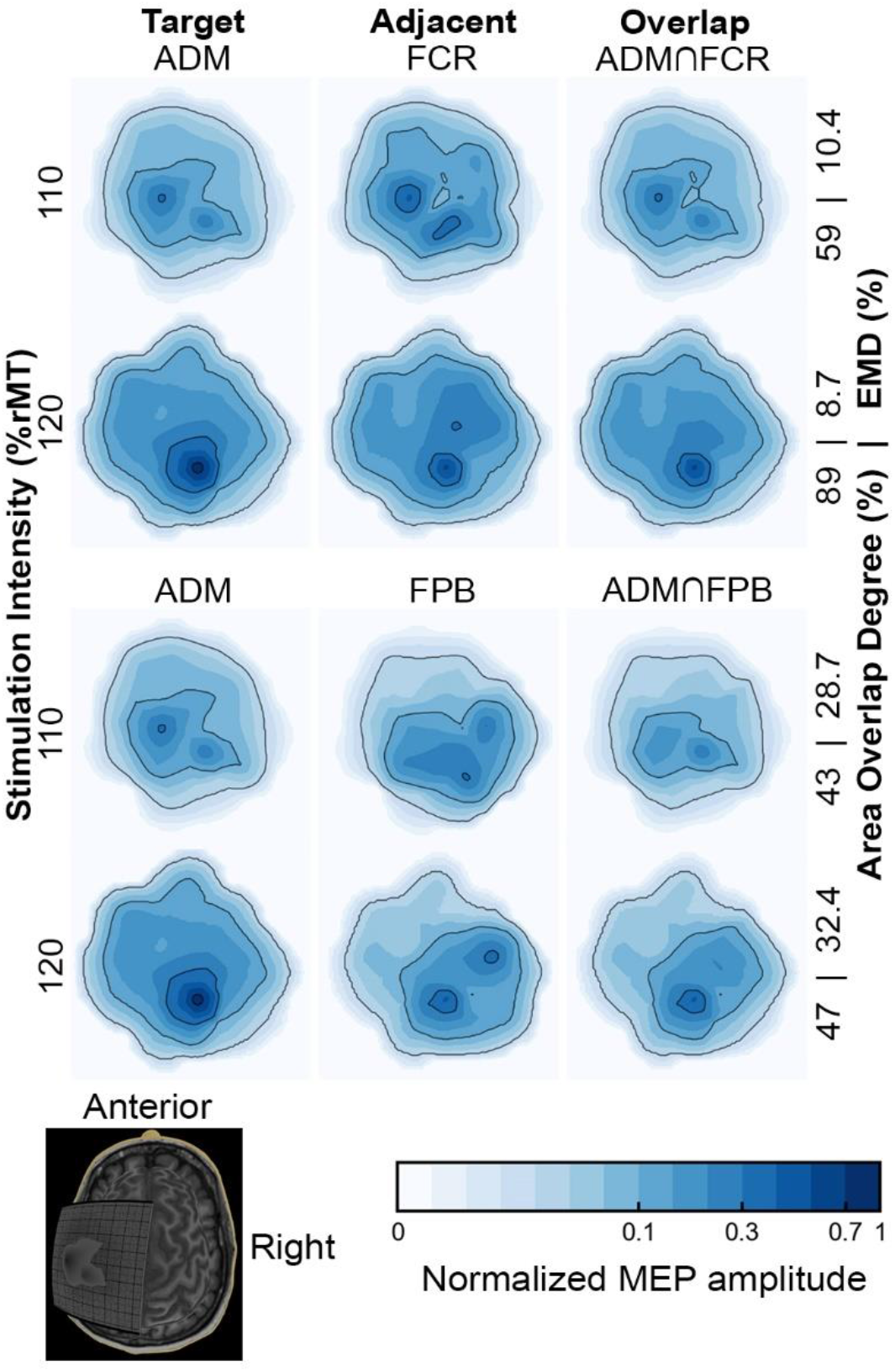
Normalized motor maps of a representative subject with low and high area and topographic similarity, and MEP amplitude affected by a higher TMS intensity. The contour line indicates the 5, 30, and 70% of the maximal MEP amplitude. Note that the map size and the area OD, but not the EMD, is significantly higher at 120% than in 110% of rMT. In addition, the increase in the area OD is greater when the EMD is lower, i.e., when the maps of the target and adjacent muscles have a more remarkable similarity. The ⋂ (intersection) symbol represents the overlap between muscles.

In turn, higher stimulation intensities are associated with stronger magnetic fields spanning a larger cortical region (van de Ruit and Grey 2016) and reflect in smoother motor maps that may lead to two outcomes. First, the higher intensity may excite neuronal populations located further from the region of interest, recruited indirectly from the excitation of intracortical neurons, thus producing progressively larger motor map areas (Schieber 2001; Nieminen et al. 2019). This stimulation leakage may overestimate the muscle representations and the region-of-interest muscle representation overlap, losing the specificity of the studied movement. Secondly, the higher intensity may delineate the full extent of the muscles’ motor representation, providing a complete picture of the muscle group coactivation. In summary, coactivation maps obtained from lower and higher stimulation intensities might provide complementary perspectives on the M1’s functional organization. The stimulation intensity needs to be carefully chosen to account for the synergy when mapping the representation of different muscles, especially in pre-surgical applications where the mapping methods must be the most accurate possible (Krieg et al. 2017).

A stimulation intensity of 120% of rMT resulted in larger representation areas of the target and adjacent muscles, corroborating previous findings (Thordstein et al. 2013; Julkunen 2014; Kallioniemi and Julkunen 2016; van de Ruit and Grey 2016). The increase in the OD is due to a higher spatial overlap between the two cortical maps than in the total area encompassed by two (or three) maps individually, as illustrated in Figure 5. Regardless of the higher overlap, the spatial distributions remain similar, and the most excitable regions of the cortex for a particular muscle seem to stay the same. Our results align with a previous study showing that an increase in stimulation intensity changes the extension of the motor representation but keeps similar topographies and centroids (van de Ruit and Grey 2016).

**Fig. 5.**
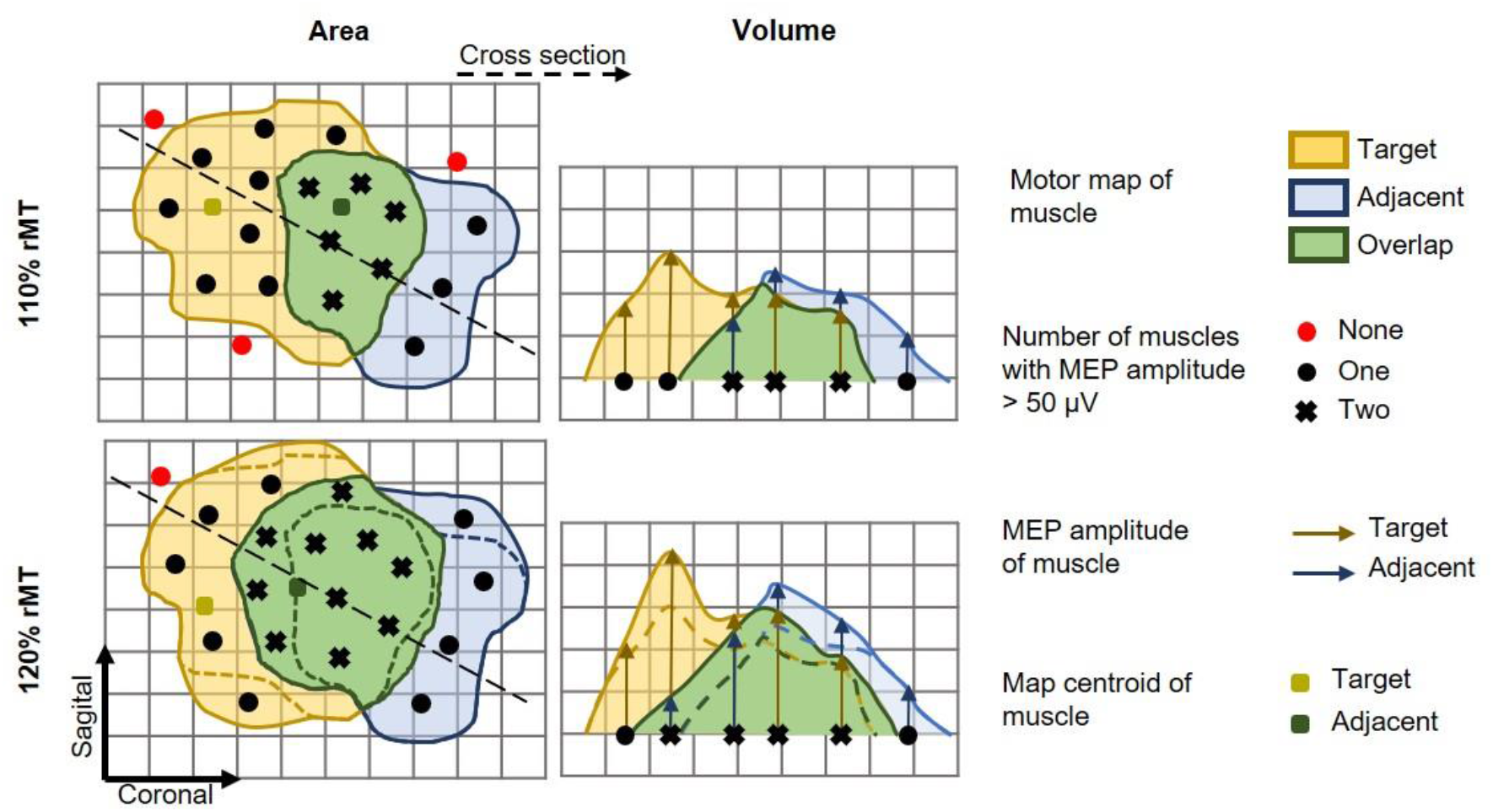
Schematic illustration of the effect of the stimulation intensity on the individual and overlap maps. In the left panel, the motor map area, and on the right, a cross-section represents the map’s height profile (MEP amplitude). Note that, at a higher stimulation intensity (120% rMT), the increase in the number of stimulation sites resulting in MEP amplitudes greater than 50 μV (black markers) from one (circle) to two (cross) muscles was more pronounced than the increase from none to one muscle. Thus, even though all maps (target and adjacent muscle and overlap) showed an increase in area, the overlap map increased more than the total map

### The cortical representation overlapping of a carpal and intrinsic hand muscles

We observed a higher overlapping between the forearm (FCR) and both intrinsic hand muscles (ADM or FPB) than between the intrinsic themselves. In addition, the centroids of the hand muscle representations are further away than those between the forearm and hand muscles. In contrast, a previous study observed higher levels of overlapping among the representations of intrinsic hand muscles when compared to those obtained between them and the carpal forearm muscles (Melgari et al. 2008; Nazarova et al. 2021). Possibly, this difference is because the subjects in our study kept their hands in a neutral posture, offering distinct proprioceptive feedback from that generated in maintaining the pronated posture (Graziano 2006), as adopted in the study by (Melgari et al. 2008). The hand posture is implicitly related to grasping and the corresponding body movements (Perez and Rothwell 2015). The influence of the hand posture on overlapping was previously associated with a dynamic modification in the neuronal network structure related to motor control (Melgari et al. 2008; Perez and Rothwell 2015; Raffin and Siebner 2019). Furthermore, the higher rMT of the forearm compared to the hand muscles may contribute to the observed higher overlap, while the low rMT of FPB may only partially stimulate adjacent muscles with higher rMT. In this sense, our results suggest that different muscles have cortical areas preferentially shared with specific muscles, as depicted in Figure 6.

**Fig. 6.**
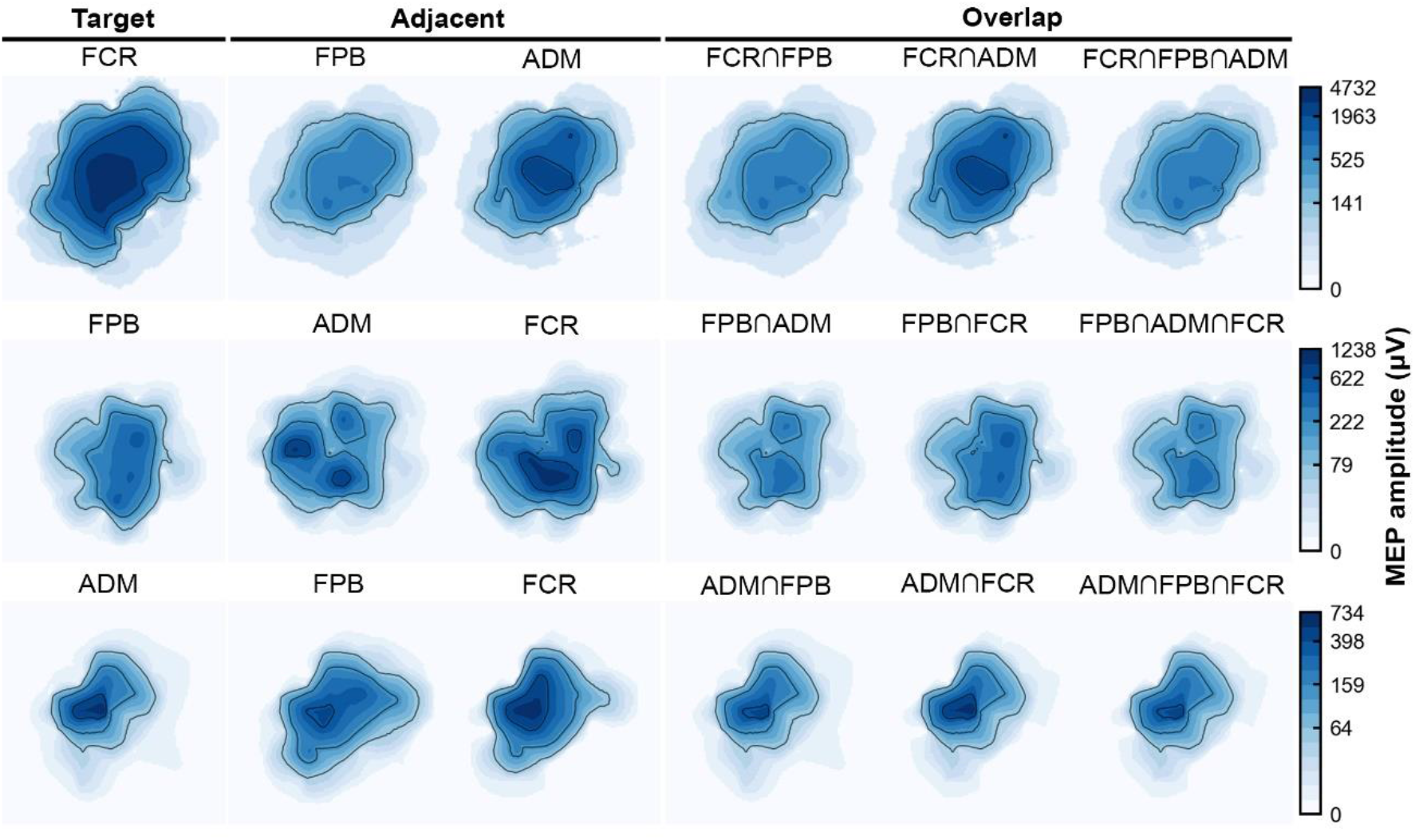
Motor maps of a representative subject at a stimulation intensity of 120% rMT, comparing topography, size, and MEP amplitudes of all possible combinations of ADM, FPB, and FCR muscle maps. The map’s anatomical references are the same as in Figure 4. The ⋂ (intersection) symbol represents the overlap between muscles.

We associate the high overlap in motor representations of upper limb muscles to a functionally organized M1 hypothesis. Our view is in agreement with previous studies that investigated the organization pattern of M1 through the nTMS motor overlapping (Wassermann et al. 1992; Wilson et al. 1993; Marconi et al. 2007; Melgari et al. 2008; Nazarova et al. 2021), and functional MRI in humans (Indovina and Sanes 2001; Leo et al. 2016). As we expected, our results revealed that the OD was smaller but still significant when both hand and forearm muscles were overlapped. This may indicate that increasing the number of overlapped motor representations reduces the overlapping degrees, resulting in greater specificity of the evoked synergies. Despite the highly overlapped representations, we have not focused on whether it reflects the ability to perform fine movements, as previous studies interpreted as indications of functional reorganization of M1 (Pearce et al. 2000; Tyc and Boyadjian 2005). However, this factor could be associated with different OD among subjects and may be tested in the future through parallel behavioral approaches.

The substantial overlapping is possibly explained by how the M1 seems to encode movements and muscle recruitment. Individual muscles appear to be recruited by a complex neuronal network instead of an individualized set of neurons, connected by a set of synergies responsible for a wide range of movements and tasks (Gentner and Classen 2006; Leo et al. 2016). Such motor representation is given by groups of functionally related neurons (Klochkov et al. 2018) following two possible mechanisms: convergence and divergence. In the convergence mechanism, a muscle has its motor representation on separate sites over M1, i.e., each site is probably overlapped with the motor representation of different groups of muscles and, thus, associated with various movements (Schieber 2001; Massé-Alarie et al. 2017). In the divergence mechanism, one site can elicit several muscles simultaneously with different intensities according to their performed movement (Schieber 2001; Melgari et al. 2008).

The divergence mechanism is observed through connections between specific pyramidal neurons and motoneurons associated with different muscles (Schieber 2001). Such a phenomenon could be related to the shared innervation between FPB with both ADM and FCR. The FPB is a hypothenar muscle composed of a superficial and a deep head that have different innervations. The deep head of the FPB is innervated by the ulnar nerve, the same innervation of the ADM. The superficial head of the FPB has the same innervation as the FCR, the median nerve (Vishram 2014). It possibly relates to the OD in the cortical representation. The action potential generated in the cortical site associated with the FPB propagates through similar pathways, resulting in a simultaneous contraction of the muscles with shared innervations. Therefore, the significant area OD probably resulted from the divergence of the overlapping motor representation indicating the synergism between the studied muscles. Animal studies further support such a view. For instance, cortical motor neurons in cats are connected to neurons of multiple muscles, not as the expected point-to-point connectivity. The synergistic interactions between neuronal populations in different cortical sites might generate descending volleys influencing various movements through the recruitment of multiple muscles (Capaday et al. 2009).

### Methodological considerations

Despite the evident extension of the overlapping representation of muscles on M1, the focality of the TMS is challenging to estimate, and stimulation can propagate over regions responsible for adjacent muscles that can contribute to a high OD (Schieber 2001; Fricke et al. 2017). However, simple models based on the coil’s center projection, like the one used in this experiment, have more than 85% accuracy and can delineate the cortical motor representation, but the stimulus propagation in the realistic cortical geometry is still disregarded (Seynaeve et al. 2019).

We should note that the EMG crosstalk in the pseudo-monopolar montage may contaminate the cortical motor representations. However, the electrode over the innervation zone might detect the direct neural drive to the muscle alleviating the crosstalk for the intrinsic hand muscles (superficial fibers running parallel to the skin) and the FCR muscle we studied (Garcia et al. 2017, 2020). Also, the cortical overlaps between the forearm and intrinsic hand muscle are more prominent than the overlap within intrinsic muscles. Thus, considering that the hand muscles are closely located and have bigger crosstalk between each other (Selvanayagam et al. 2012), our findings are significant despite this limitation and would only be further supported by reducing the potential crosstalk.

We used 100 μV MEP amplitude as a reference to estimate the rMT. The 110% and 120% of rMT stimulation intensities may correspond to slightly higher stimulator outputs when compared to protocols using 50 μV MEP amplitude. Nonetheless, our key results are the changes in motor map parameters and overlap relative to the increase in the stimulation, which are likely to occur regardless of the small deviation from the stimulator output. Moreover, the adopted protocol ensured consistent MEPs for a relatively small number of trials in each location coil location during the motor mapping procedure. We should note that only one set of muscles linked to the manual grasp movement was assessed. Our results may not generalize to muscle groups with different movement refinement, such as the lower limbs. Even so, we provide an important systemic perspective on how to evaluate the cortical motor representation.

## Conclusion

We aimed at understanding the cortical motor organization of three muscles linked to the grasping movement. Our results showed a higher cortical representation overlapping between the carpal forearm and both intrinsic hand muscles than between the intrinsic themselves. Stronger stimulation intensities led to higher overlap in the map areas but did not affect the volume and the map topographies. Our study contributes to a more detailed representation of the motor cortex associated with the functional arrangement among muscles, implying a synergistic spatial organization. Understanding the coactivation of muscle groups may provide accurate functional maps over M1. Finally, spatially accurate cortical motor mapping with nTMS can have an immediate clinical impact, for instance, when defining the eloquent brain regions during pre-surgical planning. Avoiding highly overlapped areas associated with muscular synergy would minimize deficits in the patients’ motor function (Tharin and Golby 2007; Lefaucheur and Picht 2016).

## Appendix 1

The cortical motor maps were created in the TMSmap software (Novikov et al. 2018). The software uses the coil center coordinates to fit the closest spherical surface in the least square sense. It projects them to generate a surface in a region called the patch of interest, where a quasi-regular grid is constructed. Spatial filtering is applied to merge stimulation coordinates located closer than 3 mm to compensate for the inherent errors and fluctuations of the neuronavigation system and to avoid strong influence from outliers. The mean coordinates and the median peak-to-peak MEP amplitudes of the merged coordinates are projected on the grid, and interpolation with a smoothly changing function approach is applied to construct the map. The maximal radius of the stimulation site influence on the cortical surface was set to 15 mm according to the approximate full width at half maximum (FWHM) of the electric field distribution on the cortical surface (Nieminen et al. 2015).

The area, volume, and centroid were computed from the cortical motor maps. The map area and volume were calculated according to the equations below:

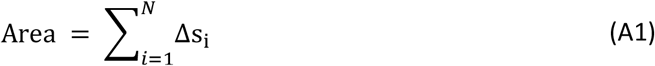

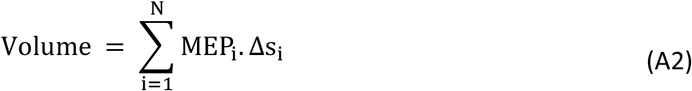

where N is the number of square grid elements with area Δs_i_ and MEP_i_ is the peak-to-peak amplitude. MEP amplitudes smaller than 50 μV were discarded from the area and volume estimates (Groppa et al. 2012). The map area represents the extent of the cortical motor representation, and the volume represents the map area weighted by the MEP amplitude. The volume is better described as an effective area. Considering a map within a particular area, the height of the map is the MEP amplitude, and it represents how strong the muscle’s response was to the stimuli applied in that area. Comparing two maps with the same area and different volumes, the extent of that muscle’s representation on M1 is equal, although the map whose height is higher has stronger muscle recruitment and representation. The centroid was calculated as:

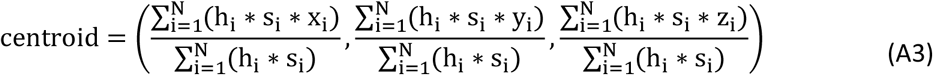

where N is the number of grid elements, h_i_ is the height of the constructed map above the grid element, s_i_ is the area and x_i_, y_i_, and z_i_ are the coordinates of the center of the grid element (Novikov et al. 2018).

The overlap map was constructed considering the area where MEP amplitudes were greater than 50 μV for all the recorded muscles. The map height is the smallest MEP amplitude across all muscles at each grid element. The area, volume, centroid and EMD for the overlap maps were calculated as described above. The size and topography parameters selected to assess the coactivation between adjacent muscles were area and volume OD, and EMD and centroids Euclidean distance, respectively, as described in Methods.

